# Widespread sex-biased gene expression reflects female-biased longevity in a species with environmental sex determination

**DOI:** 10.1101/2025.11.26.690843

**Authors:** Samantha L. Bock, Luke A. Hoekstra, Kelsi Hagerty, Robert E. Schmidt, Jessica Judson, Maxwell Adorsoo, Ritambhara Singh, Fredric J. Janzen, Anne M. Bronikowski

**Author notes:** Corresponding author: Samantha Bock. Department of Biology, Oklahoma State University, Stillwater, OK, USA. Department of Biology, University of Wisconsin – La Crosse, La Crosse, WI, USA.

## Abstract

Sexes frequently differ in life history traits including body size, lifespan, and age at sexual maturity. Aging, the progressive decline in physiological function and cellular resilience over time, is a central process contributing to sex-specific life histories, yet the mechanisms driving sex differences in aging remain largely unresolved. Long-term mark-recapture efforts revealed a striking pattern of female-biased longevity in the painted turtle (*Chrysemys picta*), a species with temperature-dependent sex determination. As a result, this species provides a compelling system to examine the mechanisms of sex-specific aging in the absence of sex chromosomes. Here, we characterize sex- and age-associated patterns in the blood transcriptomes of wild painted turtles (n = 93). We identified widespread gene expression differences between females and males (2,347 genes; 13.4% of all filtered genes). In contrast, only six genes showed significant linear relationships with continuous age in both sexes. We also employed a machine learning approach which identified distinct sets of genes for which expression was predictive of age in each sex. Age-related gene expression patterns highlight both conserved molecular pathways with known roles in aging as well as novel gene targets. These findings suggest sex-specific molecular processes underlie sex-biased demographic aging and raise questions regarding the environmental and developmental drivers of sex-biased gene expression.

## Introduction

Sex is one of the most significant sources of phenotypic variation across gonochoristic animals. Fundamental life history traits, including body size, maturation rate, reproductive strategy, and lifespan, often differ between females and males, yet the direction and magnitude of these sex differences vary widely across taxa [1,2]. Sex-associated variation in organismal aging, the progressive decline in physiological function and cellular resilience over time, is a prominent source of sex-biased life history patterns giving rise to variation in rates of demographic aging and lifespan [3]. In humans, for example, females age more slowly and consistently outlive males, a phenomenon that has persisted through environmental and social upheavals [4–6]. Long-term mark-recapture studies in wild animal populations provide further evidence for the diversity of sex-specific aging patterns, as reflected by age-associated increases in mortality and declines in fertility [1,6]. However, despite the pervasive nature of sex-specific aging patterns and critical relevance to population dynamics, we currently lack an understanding of the underlying molecular and physiological mechanisms driving these patterns and their diversity.

Hypotheses put forth to explain the mechanistic underpinnings of sex-specific life histories and aging generally invoke two processes governing biological sex differences - asymmetric inheritance and reproductive physiology [4]. While females and males largely share the same genome content, the genomic architecture of sex determination can contribute to sex-related genetic differences that alter the course of aging [7,8]. One pair of hypotheses posits that in species with sex chromosomes, the heterogametic sex (i.e., males in species with XX/XY, females in species with ZZ/ZW) should experience reduced lifespan and accelerated aging due to either the phenotypic expression of deleterious recessive alleles present on the ‘X’ (or ‘Z’) chromosome that are masked by a dominant allele in the homogametic sex (i.e., “Unguarded X” hypothesis) [9,10], or expression of deleterious genetic elements (e.g., transposable elements) on the ‘Y’ (or ‘W’) chromosome (i.e., “Toxic Y” hypothesis) [11–13]. While compelling evidence has accumulated suggesting the heterogametic sex tends to die earlier than the homogametic sex across a range of taxa [6,14,15], most of this evidence is derived from species with well-defined, heteromorphic sex chromosome systems (e.g., mammals and birds), and these hypotheses fail to explain the emergence of sex-specific aging patterns in the absence of heteromorphic sex chromosomes or genotypic sex differences altogether.

Sex-associated variation in physiology and environmental exposures likely play important roles in shaping sex-specific aging trajectories; however, disentangling the genetic, physiological, and environmental contributions to sex differences in aging has remained a daunting task, even in model organisms [1,16]. Species representing diverse modes of sex determination provide unique opportunities to examine how sex-specific aging patterns manifest under varying degrees of sex-associated genetic and environmental variation. For example, many ectotherms exhibit temperature-dependent sex determination (TSD), in which thermal signals during incubation coordinate transcriptional and physiological responses that direct the bipotential gonad to develop as an ovary or testis [17,18]. During TSD, thermal signals influence epigenetic processes to establish initial imbalances between conserved female- and male-promoting transcriptional pathways which are subsequently canalized into alternate sexual fates by downstream endocrine cues [18,19]. Interestingly, despite the absence of genetic sex differences, examples of species with TSD exhibiting differences in aging and lifespan between the sexes are abundant, particularly in the reptile clade [20,21].

One such species, the painted turtle (*Chrysemys picta*), is an emergent ecological model for testing proximate mechanisms of aging [21–23], the contributions of genes and environment to variation in aging, and the macroevolution of slow aging overall [24]. Painted turtles exhibit a pattern of TSD in which males are produced at lower incubation temperatures, females at higher incubation temperatures, and a 50:50 sex ratio at approximately 28°C [25]. In addition, long-term mark-recapture studies have revealed that males age demographically twice as fast (mortality rate doubling time: males = 10 yrs, females = 35 yrs) and live roughly half as long as females (lifespan: males = 24 yrs, females = 48 yrs) [26]. As a result, the painted turtle represents an ideal system to examine the molecular and physiological underpinnings of sex-biased aging in a wild population.

Here we take both hypothesis-driven and discovery-based approaches to understanding the drivers of sex-specific aging in painted turtles as revealed by their circulating blood transcriptomes. While studies of age-associated transcriptomic changes in wild populations are rare and primarily limited to mammals, those few studies that have documented such changes underscore the importance of studying aging phenotypes in ecologically relevant contexts [27–31]. In the present study, we examine age- and sex-associated transcriptional patterns in whole blood of wild painted turtles in a population that has been the subject of long-term mark-recapture efforts for over 30 years [32]. We sought to investigate the relative support for two hypotheses to explain observed female-biased longevity in this population. These hypotheses posit that sex-biased aging patterns are underpinned by either (1) shared age-associated transcriptomic changes that proceed at different rates in males and females, or (2) sex-specific transcriptomic patterns. We further functionally characterize sex, age, and sex-by-age associated genes to probe the specific molecular pathways contributing to expression variation and their potential links to higher order physiological aging phenotypes. Ultimately, our findings highlight potential mechanisms driving sex-biased aging patterns in a species with environmental sex determination.

## Materials and Methods

### Study Subjects and Tissue Sampling

Painted turtles (*Chrysemys picta*) were sampled from a long-term research site, Thomson Causeway Recreation Area (TCRA), in Thomson, Illinois, United States [23,33]. Field research was carried out with IACUC approval and under permits from the Illinois Department of Natural Resources. In May and June of 2017 and 2018, 95 turtles were caught with fyke nets or dip nets (N = 48 males, N = 47 females, both sexes sampled in both years). We determined the ID, sex, plastron length, and age for each animal. For individuals in this study, age was known for turtles that were first caught prior to eight years old (n = 47) based on plastron scute rings [34,35]. For all remaining turtles (n = 48), age was estimated from plastron length at first capture based on a sex-specific von Bertalanffy growth curve fit with known-age animals from the same population [21]. We drew 200 uL of whole blood from each turtle from the caudal vein within 10 minutes of capture. Immediately following blood draw, whole blood was buffered in RNAlater (Invitrogen), refrigerated at 4°C, and, within 10 days, stored at -20°C until RNA extraction and library preparation. RNA extraction was performed using the RiboPure-Blood RNA Purification Kit (Invitrogen).

### Sample Preparation and Sequencing

To avoid expected overrepresentation of globin transcripts in blood transcriptomes, we first depleted globin transcripts from the extracted RNA using oligo baits custom-designed by Lexogen (Vienna, Austria) to target the six known *C. picta* globin transcripts. Library preparation was performed by the Iowa State University DNA facility using QuantSeq 3’mRNA-Seq Library Prep Kit (Lexogen, Vienna, Austria) for Illumina. QuantSeq 3’ mRNA-Seq is an economical option for surveying the transcriptome in many samples due to reduced coverage requirements compared to traditional whole-transcript sequencing approaches. In a quantitative comparison of these methods, QuantSeq 3’ mRNA-Seq resulted in similar reproducibility [36]. Briefly, RNA was reverse transcribed using an oligo dT primer containing an Illumina-compatible sequence at the 5’ end. RNA template and globin transcripts were then degraded. The remaining cDNA was converted to dsDNA via random priming and the resulting library purified using magnetic beads. These libraries were sequenced on an Illumina HiSeq 3000 using single end 150 bp primers and forward chemistry.

### Data Processing Pipeline

Quality control, trimming, mapping, and read counting was performed according to Lexogen’s QuantSeq Data Analysis FWD parameter specifications. Read quality was assessed with FastQC (v. 0.12.1; https://www.bioinformatics.babraham.ac.uk/projects/fastqc/) [37] and MultiQC (v. 1.14) [38]. Raw reads were trimmed to remove low quality 3’ regions (average Phred quality scores < 10), poly(A) read-through, Illumina adapter contamination, and reads shorter than 20 bp using ‘bbduk.sh’ within BBMap (v. 39.19) [39]. Resulting trimmed reads were aligned to the most recent *C. picta* reference genome (ASM1138683v2; GCF_011386835.1) [40] using STAR (v. 2.7.11b) [41]. Mean alignment rate (±1 SD) was 88.4% ± 4.3. Resulting BAM files were coordinate sorted and indexed with samtools (v. 1.19.2) [42]. Gene count quantification was performed using HTSeq-count (v. 2.0.7) [43].

### Differential Gene Expression Analysis

All further analyses were conducted in R version 4.5.1 [44]. Differential gene expression analysis was performed with the DESeq2 package (v. 1.48.1) [45]. Genes were removed from analysis if the sum of counts across all samples was less than 10 or the variance of counts across all samples was less than 0.1. Of the 35,786 genes in the *C. picta* genome assembly, 17,454 genes passed filtering and were included in downstream analyses [40]. Read count normalization was performed according to DESeq2’s median of ratios method [45]. Following filtering and normalization, two samples (one female, one male) were found to have a deficit of genes with intermediate counts and were removed from the analysis.

The distribution of ages in our dataset differed between females and males according to the sex-specific age structure of the study population (Mann Whitney test p < 0.0001, n = 93; range of ages for females: 4.0 – 41.9 yrs, males: 2.4 – 19.5 yrs; **Figure 1**). To account for this in our examination of age-associated differential expression patterns, raw age was transformed via two different approaches. First, age was z-transformed within each sex to a sex-specific mean 0 and unit variance (“zAge” hereafter), allowing for the comparison of relatively old individuals to relatively young individuals across the sexes. As a complementary approach, raw age was also transformed to three categorical age groups for each sex (females yrs: G1 [4, 10), G2 [10, 16), G3 [16, 41]; males yrs: G1 [3, 6), G2 [6, 9), G3 [9, 19]) defined based on known life history transitions while attempting to distribute samples relatively evenly between groups (females: n_G1_ = 12, n_G2_ = 17, n_G3_ = 17; males: n_G1_ = 20, n_G2_ = 14, n_G3_ = 13).

**Figure 1.**
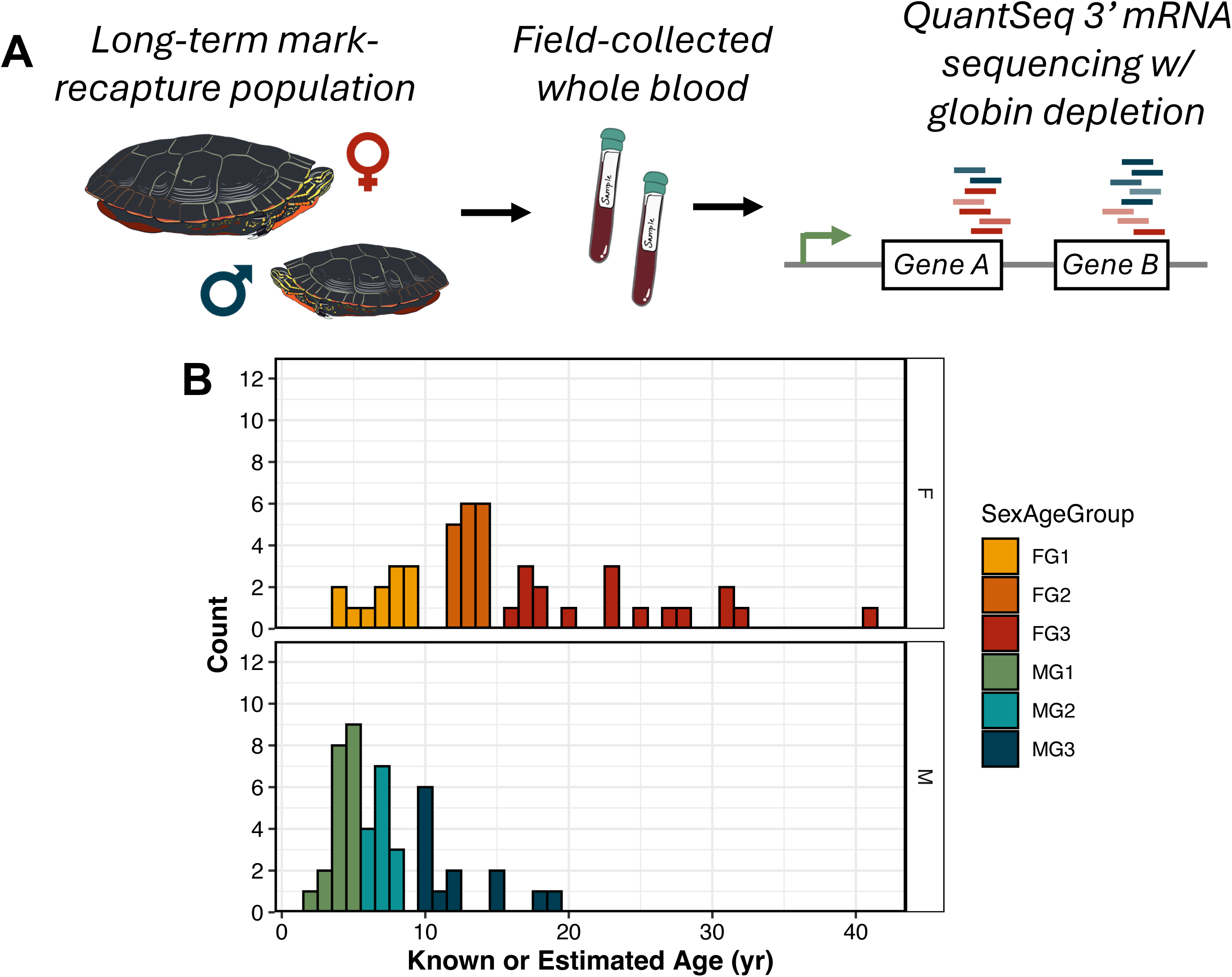
Study approach and sample distribution. (A) Adult painted turtles were caught in the field as part of long-term mark-recapture study in Thompson, IL, USA. Whole blood samples were collected in the field and preserved in RNAlater prior to extraction, library preparation, and QuantSeq 3’ mRNA sequencing. (B) Distribution of female and male samples across the lifespan reflects the sex-specific age structure of study population. Discretized age groups for each sex are depicted by bar color.

Two separate generalized linear models were fit to counts using DESeq2 [45]. The first model included an effect of sex, zAge, and their interaction (“continuous age model” hereafter). The second model included an effect of sex and categorical age group (“categorical age model” hereafter). We did not include an interaction term in the model using categorical age group based on the limitations of our sample size. DESeq2 models raw counts using a negative binomial generalized linear model with a logit link and uses sample-specific size factors as offsets. Gene-wise dispersion parameters are estimated and shrunk according to an empirical Bayes approach [45]. Prior to testing for differential expression, we removed genes exhibiting evidence of unstable coefficient estimation (‘betaIter = 100’; Supplemental Information). For outliers, Cook’s distance-based flagging was used for both models. Outlier replacement was performed for the categorical model and independent filtering based on the mean of normalized counts was enabled for both models according to DESeq2’s default settings. We used Wald tests for hypothesis testing and adjusted resulting p-values according to the Benjamini-Hochberg method. Genes were considered differentially expressed with respect to the contrast of interest according to a false discovery rate threshold (FDR) < 0.1.

### Functional Enrichment of Differentially Expressed Genes

Differentially expressed genes (DEGs) were functionally characterized according to their gene ontology terms and regulation by transcription factors. Gene ontology (GO) term overrepresentation analysis was performed using the clusterProfiler package (v. 4.16.0) [46]. To use GO annotations assigned to *C. picta* genes through the NCBI Eukaryotic Genome Annotation Pipeline, an organism-specific annotation database was constructed using the “makeOrgPackageFromNCBI()” function in the AnnotationForge package (v. 1.44.0)[47]. We then tested for overrepresentation of GO terms in each of the three annotation categories (“biological process” [BP], “cellular component” [CC], and “molecular function” [MF]) against a custom background of all filtered genes with the “enrichGO()” function in clusterProfiler. DEGs were tested separately for GO term enrichment according to the direction of log2-fold change. GO terms were considered significantly enriched according to FDR < 0.05.

DEGs were also characterized according to the transcription factors that tend to regulate them using Enrichr [48,49] and the ChIP Enrichment Analysis (ChEA) 2022 database [50]. To obtain gene names for ‘uncharacterized loci’ in the RefSeq *C. picta* genome annotation (6,192 of 17,454 filtered genes), we extracted FASTA sequences for these genes from the *C. picta* reference assembly and queried the sequences using NCBI Blastx+ (v. 2.14.1) [51]. Blastx+ was used to search for matches within the UniProt Swiss-Prot protein database [52] based on an e-value cut-off of 1 × 10^−5^. The best hit for each gene was first defined by e-value and then by percent identical matches. Hits were only retained if the percent identical matches exceeded 50%. Gene names corresponding to UniProt hits were determined using the UniProt ID mapping tool. We subsequently converted gene names to their human Ensembl alias with the g:Convert tool of the g:Profiler suite (v. e113_eg59_p19_f6a03c19) [53]. Human Ensembl aliases were used due to the availability of functional information for human genes. Genes for which a human Ensembl alias could not be assigned were excluded from subsequent enrichment analysis. Ultimately, 12,681 of 17,454 (72.6%) filtered genes were assigned gene identities corresponding to a human Ensembl gene. DEGs were tested for enrichment of transcription factor targets using Enrichr against a custom background of all filtered, annotated genes for a given contrast. A transcription factor term was considered significantly enriched according to an FDR < 0.05.

We also tested for overrepresentation of DEGs in three additional gene sets with functional significance to age-related processes – (1) candidate human age-associated genes from the GenAge database [54], (2) genes exhibiting differential expression with respect to age in human peripheral blood reported in [55], and (3) genes in the insulin/insulin-like signaling and target of rapamycin (IIS/TOR) molecular network defined in [56] (N = 61 genes). Five additional genes were included in the list of IIS/TOR network genes beyond those in [56], including *IGFLR1*, *IGFBPL1*, *IGFBP7*, *IGF2BP3*, and *IGF2BP2.* DEGs were tested for overrepresentation in these gene sets of interest using a one-sided Fisher’s exact test. Associated p-values were subjected to a Bonferroni correction within each contrast to account for repeated tests across the three gene sets of interest.

### Sex-Specific Age-Associated Feature Selection

As a complementary approach to the differential gene expression analysis, we also performed variable selection using elastic net regularized regression to identify the genes for which expression is most predictive of chronological age (in years) within each sex [57]. Prior to model fitting, raw counts for all filtered genes (n = 17,454) underwent a variance stabilizing transformation without regard for experimental group information (‘blind = TRUE’) in DESeq2. Next, the count matrix was separated by sex and subsequently z-scaled. The splitting before scaling ensured that there was no data leakage across the sex-specific matrices. The resulting normalized, scaled count matrices were modeled using an elastic net via scikit-learn, a Python package [58]. We defined a hyperparameter range of [0.001, 1] for the ‘l1_ratio’ term that balances the ridge and lasso penalties and the ‘alpha’ term, a constant that multiplies the penalty terms. We randomly sampled 100 different values in the range for the ‘l1_ratio’ and ‘alpha’ hyperparameters. A 10-fold internal cross-validation was used to identify the optimal hyperparameters that minimized mean square error [57]. Models were fitted using the entire sex-specific dataset in each iteration, and non-zero coefficients were extracted to identify selected features.

## Results

### Widespread sex-associated differential expression patterns

Females and males exhibited extensive transcriptional divergence in whole blood, with a total of 2,347 genes (13.4% of all filtered genes) exhibiting significant differential expression with respect to sex across both models according to an FDR < 0.1 (categorical age model: 2,472 sex DEGs; continuous age model: 2,693 sex DEGs; **Figure 2**; **Table S1**). Of those sex DEGs that were common across models, 1,275 genes exhibited male-biased expression and 1,072 genes exhibited female-biased expression. Among the most female-biased DEGs according to log_2_(fold change) were *ENDOU*, a gene encoding an endonuclease with key roles in immune function [59], *EGR1* and *EGR3*, which encode early growth factor proteins with roles in inflammation and regulation of cellular proliferation [60], and *AMHR2*, which encodes anti-Müllerian hormone receptor 2, a hormone receptor with key roles in reproductive function (**Figure 2A**) [61]. The DEGs exhibiting the most male-biased expression according to log_2_(fold change) included *PTGFRN*, which encodes a prostaglandin receptor inhibitor, and *COL9A2*, which encodes a component of type IX collagen (**Figure 2A**). Several other interesting candidate genes were among the most significantly male-biased in their expression, including *RAD50*, which encodes a protein involved in double-stranded DNA break repair [62], *OXCT1*, which encodes an enzyme involved in mitochondrial ketone body metabolism [63], and the homeobox gene *HOXB7*, which has also been implicated in DNA repair (**Figure 2A**) [64].

**Figure 2.**
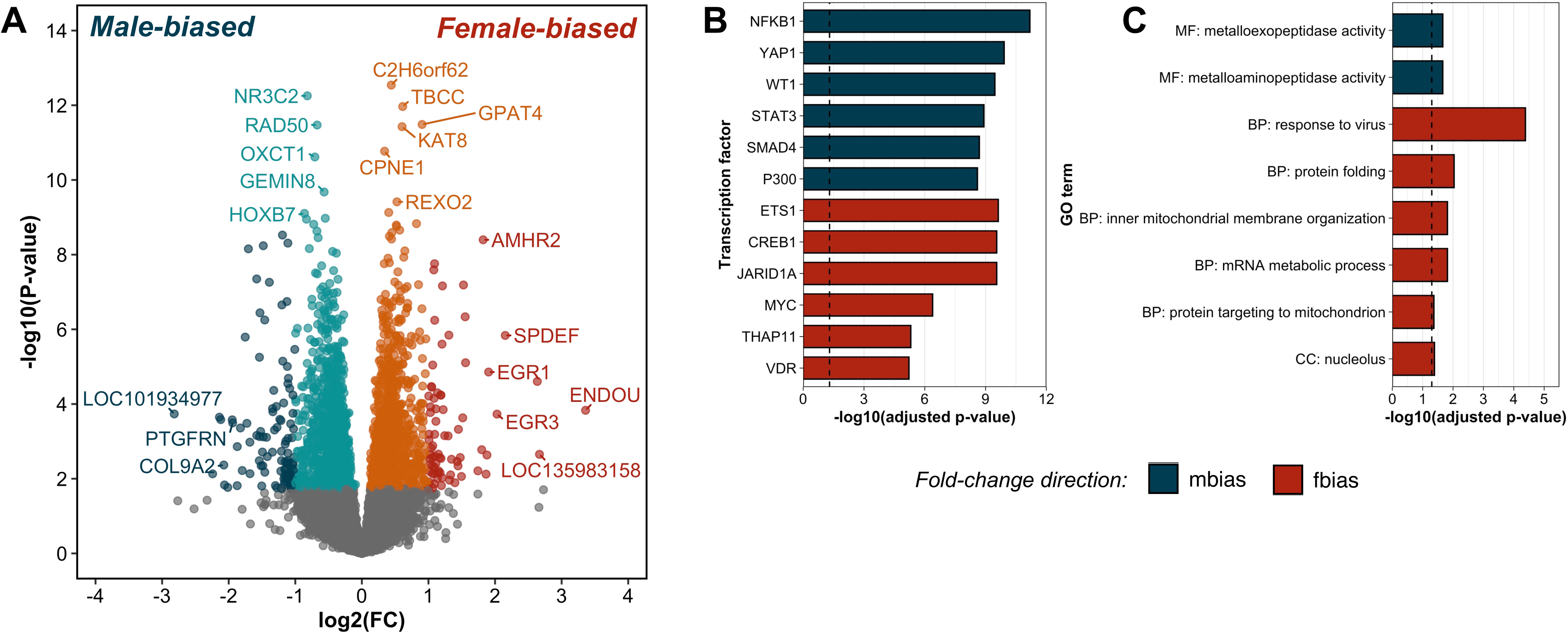
Transcriptome-wide sex differences in whole blood gene expression. (A) Volcano plot depicting the log_2_(fold change) and -log_10_(p-value) for the sex term in the model including categorical age group for each gene. Significantly differentially expressed genes (DEGs; FDR < 0.1) are depicted as colored points. Male-biased DEGs with a log_2_(fold change) > 1 are depicted by dark blue points and all other male-biased DEGs are depicted by light blue points. Female-biased DEGs with a log_2_(fold change) > 1 are depicted by red points and all other female-biased DEGs are depicted by orange points. (B) Top six most significant transcription factor regulators of male- and female-biased DEGs. (C) Subset of top enriched gene ontology terms among male-and female-biased DEGs.

### Regulation of female- and male-biased genes by distinct transcription factors

Male- and female-biased genes exhibited enrichment for regulation by distinct transcription factors with roles in immune function, cellular proliferation and apoptosis, and epigenomic modifications. In particular, the top enriched transcription factors regulating male-biased genes included nuclear factor of kappa light polypeptide gene enhancer in B-cells 1 (NFKB1), Yes-associated protein 1 (YAP1), and P300 (**Figure 2B**; **Table S2**). In contrast, the top enriched transcription factors regulating female-biased genes were ETS1, cAMP response element-binding protein 1 (CREB1), and the histone demethylase, JARID1A (**Figure 2B**; **Table S2**).

### Enrichment of molecular and physiological processes relevant to aging among sex-biased genes

Sex-biased DEGs were also enriched for distinct gene ontology terms depending on the direction of expression bias. Male-biased genes were enriched for molecular function terms “metalloexopeptidase activity” and “metalloaminopeptidase activity”, while female-biased genes were enriched for biological process terms including “response to virus”, “protein folding”, “inner mitochondrial membrane organization”, “protein targeting to the mitochondrion”, “mRNA metabolic process” and the cellular component term “nucleolus” (**Figure 2C; Table S3**). In addition, sex-associated genes collectively were enriched for members of the IIS/TOR gene network (categorical age model: odds ratio = 2.05, p_adj_ = 0.045).

### Limited scope of age-associated gene expression

Age-associated gene expression patterns were much more limited relative to sex-associated patterns (**Figure 3**). Only three genes (*IQGAP3*, *TOP2A, LOC101942607 –* zinc-finger protein RFP-like) were significantly differentially expressed between the youngest (‘G1’) and middle-aged individuals (‘G2’; **Figure 3A**) and only seven genes were significantly differentially expressed between the youngest (‘G1’) and oldest individuals (‘G3’; **Figure 3B**; **Figure S1**; **Table S4**). Six genes showed significant linear relationships with continuous age, half of which showed greater expression at older ages and half of which showed greater expression at younger ages (**Figure 3C**; **Figure 4A**; **Figure S2**; **Table S4**). For the purposes of exploratory enrichment analysis, we loosened the significance threshold to include age-associated genes with unadjusted p-values < 0.1. This analysis revealed functional differences between genes exhibiting positive and negative associations with age. Among genes with expression related to continuous age according to the loosened significance threshold, genes showing greater expression in younger individuals (i.e., negative age-association) were enriched for regulation by transcription factors including KDM2B, FOXO1, MAF, and TCF7 (**Figure 4B**; **Table S5**). Interestingly, *TCF7*, which encodes a transcription factor involved in T cell differentiation [65], was also the only gene that showed significant differential expression with respect to both continuous age and categorical age group (**Figure 3**). Genes with negative age-associated expression were also enriched for gene ontology terms primarily related to immune function, including the biological process terms “regulation of immune system process”, “leukocyte activation”, and “antigen processing and presentation” (**Figure 4C**; **Table S6**). In contrast, genes showing greater expression in older individuals were enriched for regulation by MYC, YY1, and MYCN and were enriched for gene ontology terms related primarily to transcriptional and translational regulation, including the terms “amide biosynthetic process”, “tRNA aminoacylation”, “mRNA splicing via spliceosome”, and “ribonucleoprotein complex” (**Figure 4C**; **Table S6**).While age-associated DEGs were not enriched for human age-related genes in the GenAge database, age-associated DEGs did exhibit enrichment (zAge: odds ratio: 1.46; p_adj_ = 2.33E-05; G1vG3: odds ratio: 1.38; p_adj_ = 0.0003) for genes showing age-associated expression in human peripheral blood [55]. These genes did not necessarily exhibit concordant directions in their relationship with age between turtles and humans. In fact, of the 191 overlapping genes, 97 (50.7%) exhibited concordant patterns of age-association and 94 exhibited discordant patterns of age-association across species.

**Figure 3.**
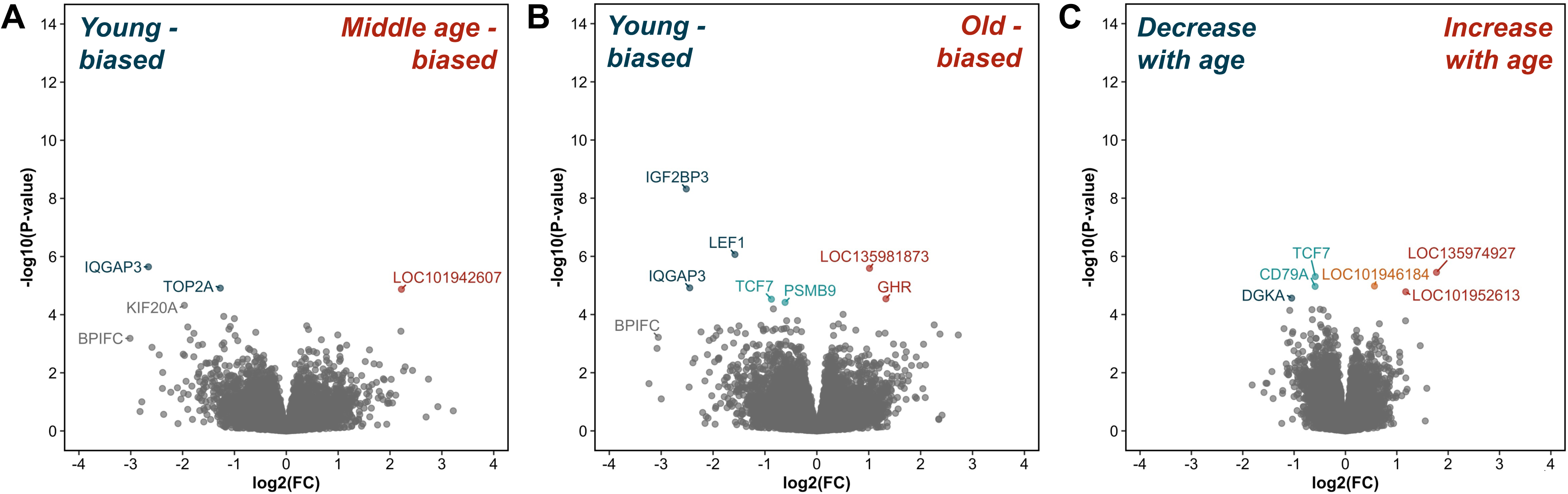
Gene expression patterns associated with categorical age group and continuous age. Volcano plot depicting the log_2_(fold change) and -log_10_(p-value) for the (A) young (G1) versus middle-age (G2) contrast, (B) young (G1) versus old (G3) contrast, and (C) z-scaled age term in the models including either categorical age group or continuous age for each gene. Differentially expressed genes (DEGs) showing greater expression in younger animals are depicted by dark blue points if log_2_(fold change) > 1 and light blue points otherwise. DEGs showing greater expression in older animals are depicted by red points if log_2_(fold change) > 1 and orange points otherwise.

**Figure 4.**
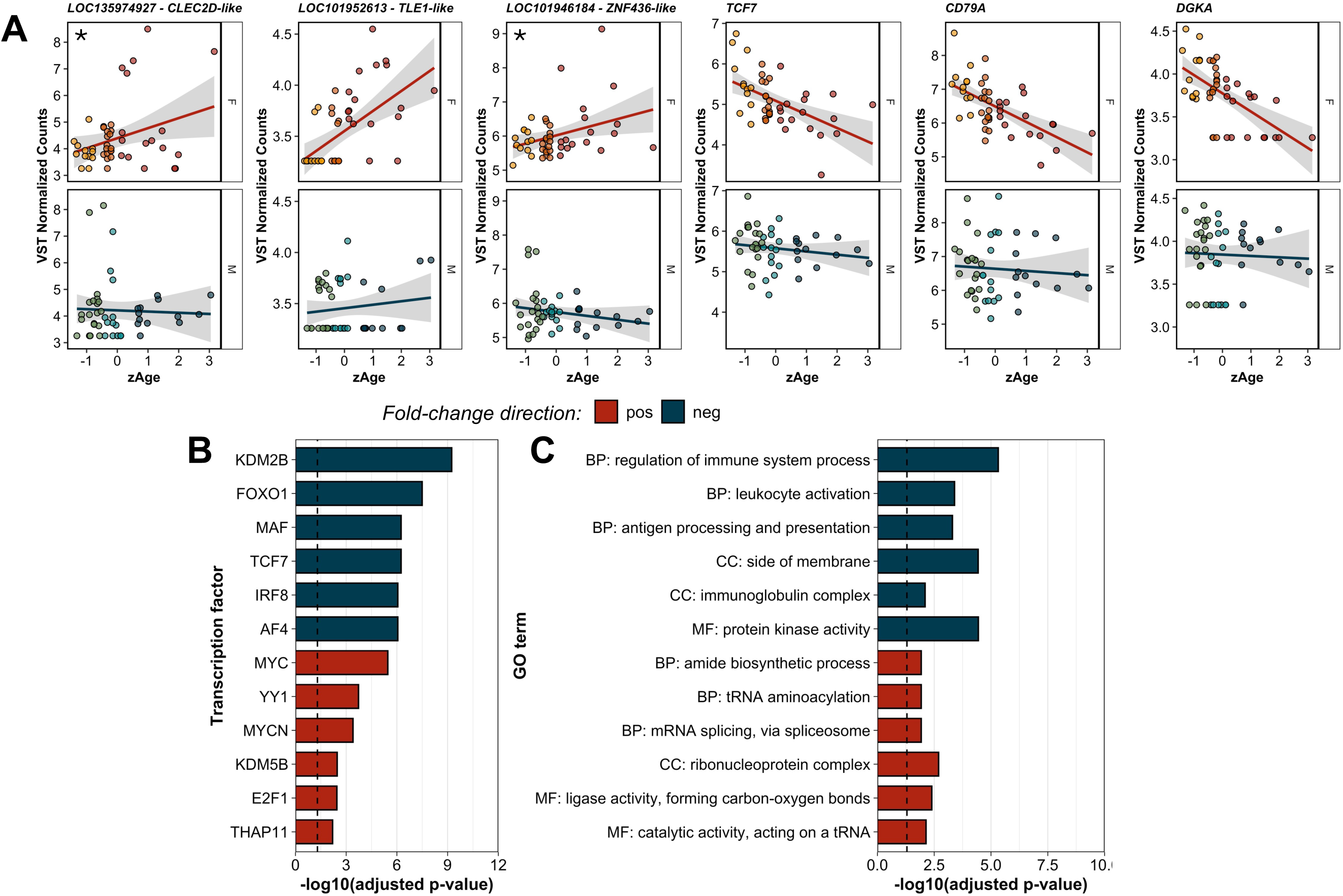
Linear age-associated gene expression patterns. (A) Variance stabilized counts across z-scaled age in males and females for each of six genes exhibiting significant differential expression with respect to continuous age. Asterisks indicate genes for which there was a significant sex-by-age interactive effect on expression. (B) Top six most significant transcription factor regulators of genes with expression positively and negatively related to continuous age according to unadjusted p-value < 0.1. (C) Subset of top enriched gene ontology terms for genes with expression related to continuous age according to unadjusted p-value < 0.1.

### Sex-by-age interactive effects on expression

Only two genes (*LOC135974927*, *LOC101946184*) that were significantly related to continuous age exhibited a significant sex-by-age interactive effect on expression according to FDR < 0.1 (**Figure 4**). While both these genes are currently uncharacterized in the *C. picta* genome annotation, they are predicted to encode C-type lectin domain family 2 member D (CLEC2D)-like and zinc finger protein 436 (ZNF436)-like proteins, respectively, based on sequence similarity. For both genes, expression increased with age in females but did not change or decreased slightly with age in males (**Figure 4**).

### Sex-specific transcriptional markers of age

Next, we used the sex-specific predictive models to extract the genes with the highest non-zero coefficients when predicting chronological age. The female-specific model yielded many more genes (300+ genes) with non-zero coefficient values compared to the male-specific model (39 genes). Only 3 common genes were assigned non-zero coefficients in both female and male models - *IGF2BP3, LOC101953725, LOC135972840*. We selected a percentile threshold based on the distribution of the values of the non-zero coefficients. For the female-specific genes, we analyzed top 32 genes (90-th percentile), and for the male-specific predictions, we looked at the top 10 genes (75-th percentile). Furthermore, these top genes for each sex-specific model were fully distinct (**Figure 5, Table S7**). This result follows similar patterns to the DEG analysis, where we observed an extensive transcriptional divergence.

**Figure 5.**
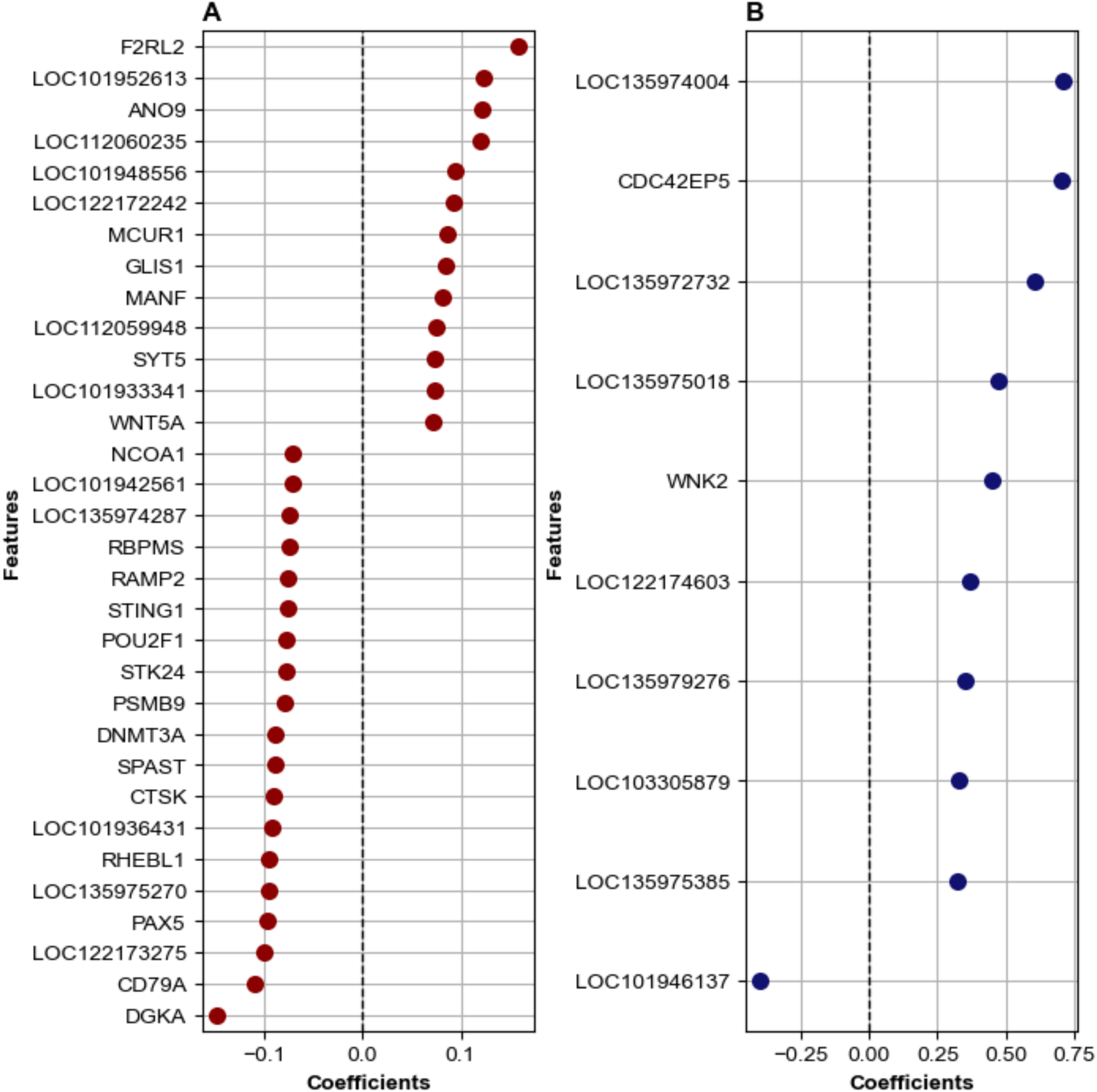
Top gene expression predictors of age in females and males. Non-zero coefficient estimates for genes included in at least 90-th percentile for females (A) and at least 75-th percentile for males (B) when elastic net regularized regression was implemented to predict chronological age in each sex.

## Discussion

Long-term mark-recapture data in a wild population of painted turtles suggests females exhibit slower demographic aging and substantially greater longevity than males [23,26], but the molecular and physiological mechanisms underlying this pattern are unknown. In the present study, we hypothesized that sex-biased aging may be reflected by (1) shared age-associated gene expression changes that proceed at faster rates in the faster aging sex (i.e., males) or (2) sex-specific gene expression patterns. Our results demonstrate that the transcriptomes of blood cells in wild adult painted turtles exhibit widespread sex-associated variation across the life course. In contrast, we did not find evidence for more rapid age-associated transcriptional changes in males. In fact, age-associated transcriptional patterns were relatively limited. Fewer than ten genes exhibited an association between age and expression, regardless of whether age was treated as a continuous or categorical trait, compared to over one thousand genes differing by sex. These results stand in sharp contrast to gene expression patterns observed in the whole blood of free-living mammals, where sex differences are limited in scope and age drives expression variation [27,31]. This discrepancy is especially interesting considering differences in the sex determination systems of these taxa. In mammals, sex is determined at conception by heteromorphic sex chromosomes, while in painted turtles, sex is determined by temperature [66]. Accordingly, sex-biased gene expression in painted turtles must result from sex-associated transcriptional regulation, while in mammals, it may also result from sex-limited genes located on the Y chromosome or genes escaping dosage compensation on the X chromosome [16,67]. One might predict that mammals would show more pronounced and widespread sex-biased gene expression in a somatic tissue. Yet, our findings run counter to this prediction. Reconciling this discrepancy necessitates further investigation into the regulatory mechanisms driving somatic sex-biased gene expression in the absence of sex chromosomes.

Transcriptional patterns observed in whole blood may reflect changes in gene expression or changes in populations of constituent cell types (e.g., monocytes, lymphocytes, and granulocytes). In turtles and other nonmammalian vertebrates, erythrocytes are transcriptionally active [68] and previous work in the same study population observed no significant effects of age and only minor effects of sex on cell proportions (**Table S8**). Taken together, our results are likely driven primarily by sex differences in gene expression rather than sex differences in immune cell composition. In line with the observation that sex contributes to broad transcriptional differences in whole blood beyond its influence on immune composition, a study of whole blood transcriptomes in the Mojave desert tortoise (*Gopherus agassizii*), another species with TSD, showed that sex-biased genes outnumbered genes responding to bacterial pathogen (*Mycoplasma agassizii*) infection by an order of magnitude (n = 40 genes associated with bacterial treatment, n = 1,037 genes associated with sex)[69]. However, the degree to which sex shapes blood cell type-specific transcriptional patterns warrants further inquiry, as studies in mammals suggest an important role for cell type-specific gene expression in contributing to sex-biased disease susceptibility [70].

The sex-biased gene expression we observe likely results from a combination of three non-exclusive drivers – (1) regulation by sex steroid hormones, (2) responses to sex-specific environmental exposures, and (3) lasting influences of the incubation environment. Sex steroid hormones regulate sex-biased gene expression across tissues and interact with the immune system [71,72]. Regulation of gene expression by sex steroid hormones depends on circulating concentrations of sex steroids, steroid binding proteins, and hormone receptor expression, all of which exhibit strong temporal fluctuations in seasonal breeders [73,74]. The timing of peaks in circulating sex steroids overlaps, at least in part, with our sampling period, where adult turtles were captured between May and June [73,74]. Indeed, genes exhibiting male-biased expression were enriched for targets of the androgen receptor (Table S2), while female-biased genes were not. It is important to consider that individual circulating sex steroid levels depend on the reproductive status of an individual at the time of sampling [73,74]. Adult female painted turtles may not reproduce every year [75], and females included in our study likely vary in their individual reproductive status. Further studies measuring plasma concentrations of steroid hormones in parallel with blood cell gene expression in the same individuals are needed to clarify the roles of hormones in regulating sex-biased gene expression. In addition, repeated sampling across different times of the year could reveal the degree to which sex-biased gene expression fluctuates by season.

The transcriptome changes dynamically with shifts in extrinsic (e.g., temperature, water availability, predator presence, pathogen exposure) and intrinsic factors (e.g., physiological state). Environmental factors associated with sex differences in behavior and microhabitat use during the nesting season likely contributed to the sex-biased gene expression we observe. Sex-specific microhabitat use and movement behavior is well-documented in this species [76,77]. Beyond female-specific terrestrial movement associated with nesting, females also exhibit greater basking behavior [76] and greater variability in body temperature than males during the nesting season [78]. Enrichment of sex-biased genes for metabolic and mitochondrial processes point to a potential role of temperature in shaping observed transcriptional differences. However, this role is difficult to parse from other co-occurring environmental and intrinsic factors. While numerous studies document widespread transcriptomic responses to temperature in ectotherms [79], rarely are these studies conducted in the context of the environmental complexity present in nature. To address this knowledge gap, future investigations could benefit from measuring body temperatures of free-living ectotherms at the time of sampling and employing paired studies of sex-biased gene expression in controlled and natural conditions to determine the extent to which sex-biased patterns persist in the absence of sex-specific environmental exposures.

In addition to potential influences of current environmental conditions, sex-associated transcriptional patterns may also result from lasting influences of the incubation environment. In painted turtles, a relatively narrow range of incubation temperatures produces mixed-sex clutches (i.e., transitional range of temperatures: 1.0-3.4°C) [25,33]. As a result, female and male painted turtles often differ in the thermal environment experienced during incubation. The degree to which incubation temperature exerts lasting effects on adult gene expression in somatic tissues is largely unknown. A few studies demonstrate persistent signatures of incubation temperature on molecular phenotypes (e.g., gene expression, DNA methylation) in juvenile or newly mature adult ectotherms [80,81], however little is known regarding how persistent these effects are throughout the life course. Even less is known about how developmental environments shape adult sex differences and subsequent aging phenotypes [82,83]. Given the observed enrichment of sex-biased genes for regulation by chromatin modifiers, it is interesting to consider the ontogeny of sex-biased gene expression and how early environmental signals may be integrated into epigenomic changes that shape adult sex-specific life history trajectories.

Age-associated transcriptional patterns, in contrast to sex-associated patterns, were limited to a small number of genes. However, those genes that did exhibit a significant association with age in adult turtles highlight potentially conserved processes contributing to aging across amniotes. For example, *TCF7*, which encodes a transcription factor regulating T-cell stemness and regenerative potential [84], and *LEF1*, which encodes a transcription factor regulating age-associated gene expression in human and mouse immune cells [85], were among genes exhibiting increased expression in young turtles. Interestingly, genes with greater expression at younger ages were also enriched for targets of *TCF7*. Recent evidence has identified maintenance of *TCF7* expression with age as a key contributor to immune resilience and healthspan in a large human cohort [86]. Other genes exhibiting age-associated expression in painted turtles that have been implicated in human aging or age-related pathologies (i.e., cancer) include *TOP2A*, encoding a protein involved in the alternate lengthening of telomeres pathway [87], and *IGF2BP3*, encoding an mRNA binding protein implicated in human tumorigenesis [88]. These findings, along with the observation that genes with age-associated expression in human peripheral blood were enriched among age-associated genes in our study [55], lends further evidence to the hypothesis that a subset of shared mechanisms (e.g., insulin/insulin-like signaling and target of rapamycin) contribute to aging in distantly-related species with divergent life history trajectories [89]. At the same time, genes in which the direction of age-association varies between turtles and humans, as well as novel turtle-specific age-associated genes, warrant attention as targets for future comparative studies unraveling underpinnings of longevity.

In addition to only two genes exhibiting significant interactive effects of age and sex on expression, genes for which expression was most predictive of age in each sex were largely distinct, suggesting a role for sex-specific processes in contributing to observed sex-biased demographic aging. However, deciphering the genes with causal links to higher order aging phenotypes and the mechanisms driving their sex-specific expression will require investigations into the functions of these genes and their regulation in a broader comparative context. Even so, our machine learning approach uncovered novel candidate genes contributing to sex-biased aging that were not revealed by our differential gene expression analysis (e.g., *CDC42EP5*, *WNK2*, *PAX5*, *RHEBL1*). This is especially interesting given the rapid proliferation of molecular age predictors built using machine learning algorithms to identify informative genomic regions or genes from large epigenetic or transcriptional datasets (e.g., [90–92]). Our results underscore how such approaches represent a complementary way of discovering candidate molecular processes contributing to intraspecific differences in longevity.

In the present study, we describe broad sex-associated transcriptional differences in the whole blood of painted turtles, raising new hypotheses about the drivers of sex-specific life history trajectories in this species. Our findings highlight fundamental questions about how sex-biased aging and life history patterns emerge from a shared genome. Ectothermic tetrapods offer a particularly insightful context to address these questions as they exhibit both variable aging rates and tremendous diversity in modes of sex determination [24,93]. For example, multiple reptile and amphibian species possess genotypic sex determination that can be overridden by certain environmental conditions [94–96]. Investigating sex- and age-associated phenotypes in naturally occurring sex reversed individuals represents an exciting future direction to further disentangle the genetic, physiological, and environmental drivers of sex-biased life history patterns in wild populations.

## Acknowledgements

The authors would like to thank the numerous students and colleagues who participated in the fieldwork associated with this project. The authors would also like to thank the Iowa State University DNA Facility for its support in generating the sequencing data. This work was made possible through support from the National Science Foundation (NSF DBI-2213824 to AMB and RS, NSF IOS-1257857 to FJJ) and the National Institute on Aging (NIH/NIA R01AG049416 to AMB and FJJ).

## Conflict of Interest

We declare that we have no competing interests.

## Data Availability

Upon acceptance, raw sequencing reads and associated metadata will be deposited in the NCBI Gene Expression Omnibus (GEO) database. Scripts used in data analysis will be publicly available on Github.

## Author Contributions

*Conceptualization:* FJJ, AMB; *Data curation:* SLB, LAH; *Formal analysis:* SLB, MA, RS; *Investigation:* SLB, LAH, KH, RES, JJ, MA, RS, FJJ, AMB; *Funding acquisition:* FJJ, RS, AMB; *Supervision:* FJJ, AMB; *Writing – original draft:* SLB, MA, RS; *Writing – review & editing:* SLB, LAH, KH, RES, JJ, MA, RS, FJJ, AMB.

---

**Figure S1.**
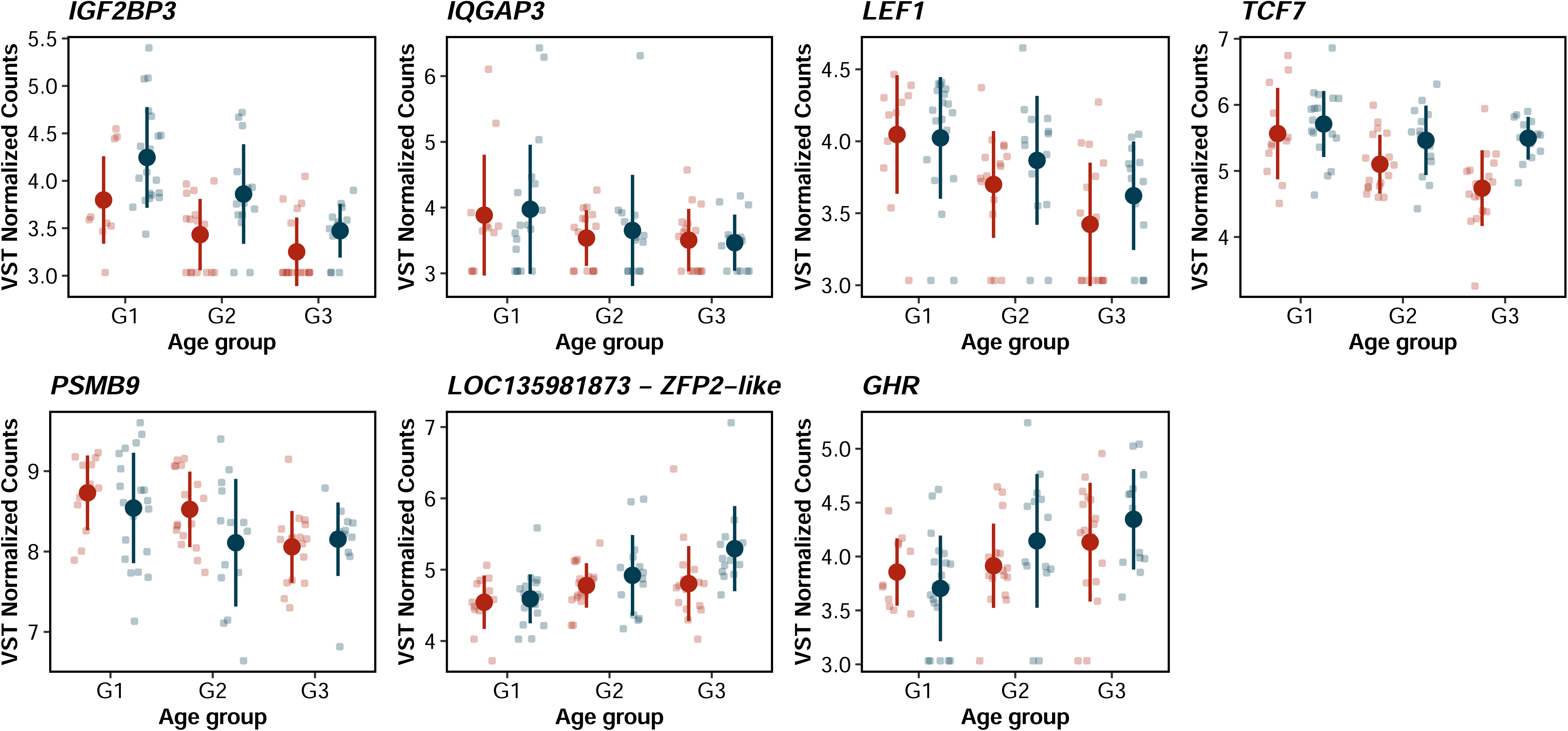
Variance stabilized counts across categorical age groups for the seven genes exhibiting significant differential expression between young (G1) and old (G3) individuals.

**Figure S2.**
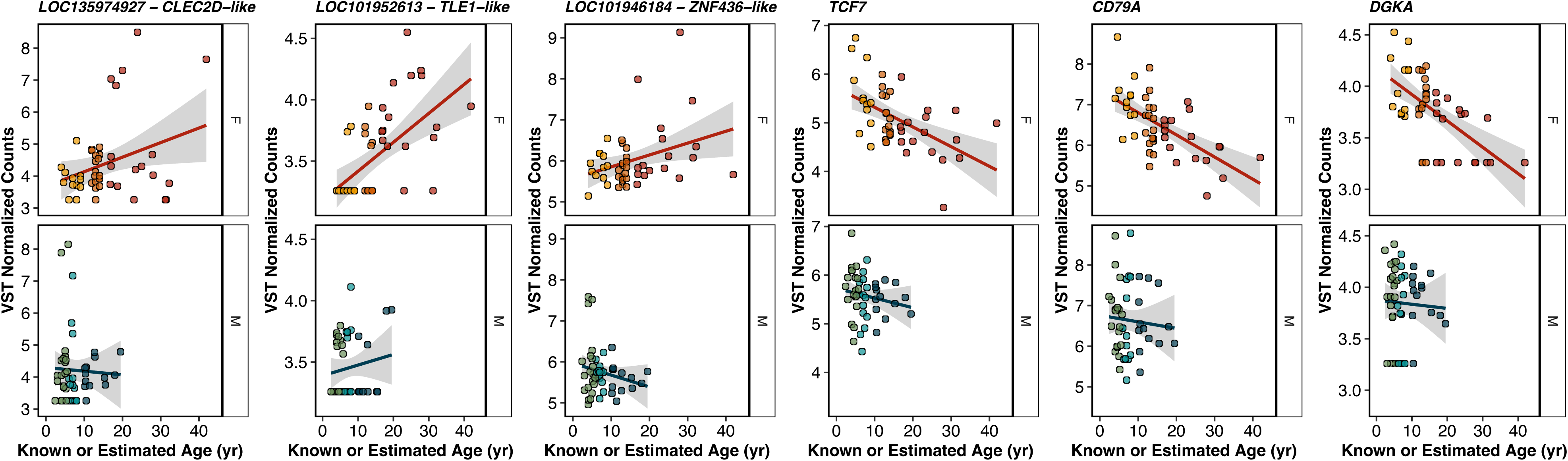
Variance-stabilized counts across raw age (yrs; unscaled) in males and females for each of six genes exhibiting significant differential expression with respect to continuous age.

## Notes

### Competing Interest Statement

The authors have declared no competing interest.

